# DcPAR1 is Required for Development and Carotenoid Synthesis in the Dark-Grown Carrot Taproot

**DOI:** 10.1101/2021.09.22.461395

**Authors:** Daniela Arias, Angélica Ortega, Christian González, Luis Felipe Quiroz, Jordi Moreno-Romero, Jaime F. Martínez-García, Claudia Stange

**Author notes:** **Author contributions**: C.S. conceived the research plans; C.S. and J.F.M-G. supervised the experiments; D.A. and A.A performed the experiments; J. M-R, C.G and L.F.Q. performed some experiments with D.A; J.F.M-G. provided technical assistance to D.A.; C.S. designed the experiments and analyzed the data; C.S. conceived the project and wrote the article with contributions of all the authors; C.S. agrees to serve as the author responsible for contact and ensures communication. **Funding:** Chilean ANID Fondecyt Grant 1180747 (CS), Spanish MINECO- FEDER grant BIO2017-85316-R to JFM-G, European Commission contract H2020-MCSA-IF-2017 (proposal 797473) to JM-R.

## Abstract

Light stimulates carotenoid synthesis in plants during photomorphogenesis through the expression of *PHYTOENE SYNTHASE* (*PSY*), a key gene in carotenoid biosynthesis. The orange *Daucus carota* (carrot) synthesizes and accumulates high amounts of carotenoids in the taproot that grows underground. Contrary to other organs, light impairs carrot taproot development and represses the expression of carotenogenic genes such as *DcPSY1* and *DcPSY2* reducing carotenoid accumulation. By means of an RNA-seq, in previous analysis we observed that carrot *PHYTOCHROME RAPIDLY REGULATED 1* (*DcPAR1*) is more expressed in the underground grown taproot respect to those grown in light. PAR1 is a transcriptional cofactor with a negative role in the shade avoidance syndrome regulation in *Arabidopsis thaliana* through the dimerization with PHYTOCHROME INTERACTING FACTORs (PIFs), allowing a moderate synthesis of carotenoids. Here we show that overexpressing *AtPAR1* in carrot produces an increment of carotenoids in taproots grown underground as well as higher *DcPSY1* expression. The high identity of AtPAR1 and DcPAR1 let us to suggest a functional role of DcPAR1 that was verified through the *in vivo* binding to AtPIF7 and the overexpression in *Arabidopsis*, where it increments *AtPSY* expression and carotenoid accumulation together with a photomorphogenic phenotype. Finally, *DcPAR1* antisense carrot lines presented a dramatic decrease in carotenoids levels and in the relative expression of key carotenogenic genes as well as impairment in taproot development. These results let us to propose that DcPAR1 is a key factor for secondary root development, plastid differentiation and carotenoid synthesis in carrot taproot grown underground.

**One-sentence summary:** DcPAR1 is a key factor for secondary root development, plastid differentiation and carotenoid synthesis in carrot taproot grown underground.

## Introduction

Light is an essential cue for plant development and growth since it is required for photosynthesis and photomorphogenesis (Tripathi et al., 2019). In both processes, the plastidial pigments chlorophylls and carotenoids fulfil crucial roles. Synthesis of chlorophylls and carotenoids is strongly promoted by light (Stange and Flores, 2012). Carotenoids also provide yellow, orange and red colors to flowers, fruits and some roots. These pigments are not only powerful antioxidant that protect plants from photooxidative damage but also precursors of important compounds such as the phytohormones strigolactones and abscisic acid (ABA) that affect plant development (Hirschberg, 2001; Howitt and Pogson, 2006; Sandmann et al., 2006; Rodríguez-Concepción, 2010; Ruyter-Spira et al., 2013; Sandmann, 2015; Simpson et al., 2016a). The first enzyme in carotenoid synthesis and the most regulated at the transcriptional and posttranscriptional level is phytoene synthase (PSY) (Maass et al., 2009; Rodríguez-Villalón et al., 2009; Rosas-Saavedra and Stange, 2016). Whereas in *Arabidopsis thaliana* this enzyme is encoded by the *AtPSY* gene, in carrots (*Daucus carota*) it is encoded by two genes, *DcPSY1* and *DcPSY2*.

When seedlings germinate in the dark, they exhibit a pale yellow or rather whitish color due to the lack of pigmentation (etiolated seedlings). In these conditions, PHYTOCHROME INTERACTING FACTORs (PIFs) accumulate and repress photomorphogenesis. PIFs are a subfamily of basic helix–loop–helix (bHLH) transcription factors that act as central mediators in a variety of light-mediated responses, including transitions from etiolated to de-etiolated seedlings and plants responses to vegetation proximity (Leivar et al., 2009; Pham et al., 2018). As part of their repressive role of the photomorphogenic development in the dark, PIFs bind to *AtPSY* Light Responsive Elements (LREs, such as G-box), preventing their expression and impeding the synthesis and accumulation of carotenoids in these conditions (Toledo-Ortiz et al., 2010). Once the emerging seedlings perceive the light, photomorphogenesis is triggered (de-etiolation), which results in inhibition of stem elongation (e.g., hypocotyls or epicotyls), promotion of cotyledon or leaf expansion and promotion of chlorophyll and carotenoids synthesis, which result in the acquisition of the characteristic green color of plants (Quail, 2002; Kami et al., 2010; De Wit et al., 2016). Light perception is carried out by photoreceptors such as phytochromes (PHYs), cryptochromes (CRYs) and phototropins (Kami et al., 2010; Stange and Flores, 2012; De Wit et al., 2016; Quian-Ulloa and Stange 2021). In particular, PHYs participate in the induction of the expression of key carotenogenic genes such as *PSY* (Hirschberg, 2001; Bramley, 2002; Simkin et al., 2003; Woitsch and Römer, 2003; Adams-Phillips et al., 2004; Giovannoni, 2004; Rodriguez-Concepcion and Stange, 2013). During de-etiolation, the light activation of PHYs triggers PIF phosphorylation, which are subsequently degraded by the proteasome and/or inactivated, therefore making them unable to bind to LREs (Bae and Choi, 2008; Shen et al., 2008; Shin et al., 2009; Toledo-Ortiz et al., 2010). This results in a rapid de-repression of *PSY* gene expression and a burst in the production of carotenoids in coordination with chlorophyll biosynthesis and chloroplast development for an optimal transition to photosynthetic metabolism (Toledo-Ortiz et al., 2010). In addition to repressing gene expression, PIFs also promote the expression of dozens of genes in the dark that are also rapidly down-regulated after seedling de-etiolation, such as *A. thaliana PHYTOCHROME RAPIDLY REGULATED 1* (*AtPAR1*), that encodes for a transcriptional cofactor of the bHLH family. AtPAR1 physically interacts with PIFs, thus partially preventing the binding of PIFs to LREs of photomorphogenic genes hence producing intermediate phenotypes between dark- and light-grown plants regard pigment synthesis and plant lenght (Bou-Torrent et al., 2015). *A. thaliana* lines that overexpress *AtPAR1* present an increase in total carotenoids in photosynthetic tissues, an increment in *AtPSY* expression and an exacerbated photomorphogenic phenotype shown as a reduced hypocotyl length (Roig-Villanova et al., 2007; Hao et al., 2012; Zhou et al., 2014; Bou-Torrent et al., 2015).

Unlike other plants, orange carrots accumulate large amounts of carotenoids in its underground and dark-grown taproots (Stange et al., 2008; Fuentes et al., 2012; Rodriguez-Concepción and Stange, 2013). Surprisingly, root exposure to white light (W) causes a reduction in the expression of carotenogenic genes such as *PSYs* and *LYCOPENE BETA-CYCLASES* (*LCYBs*) and a decrease in carotenoid levels (Fuentes et al., 2012; Llorente et al., 2017; Arias et al., 2020). Indeed, the carrot root grown in W has a thinner and greener phenotype and presents an enrichment of chloroplasts instead of carotenoid accumulating chromoplasts (Fuentes et al., 2012). The expression profile between carrot taproot grown in W and underground (dark-grown) was achieved and compared through an RNA-Seq (Arias et al., 2020). This led us to identify several carrot genes related to photomorphogenesis, such as *DcPHYA*, *DcPHYB*, and *DcPAR1*, whose transcripts accumulated in the underground dark-grown taproot but was rapidly reduced in the root when exposed to W (Arias et al., 2020).

In this work we aimed to functionally characterize *DcPAR1* to determine its role in photomorphogenesis and in carotenoid synthesis in Arabidopsis and carrot. We showed that overexpression of *AtPAR1* in transgenic carrot present higher levels of carotenoids in taproots as well as higher *DcPSY1* expression suggesting a key role of the *PAR1* in carrot carotenoid synthesis. The high similarity of AtPAR1 and DcPAR1 let us to suggest a functional role of DcPAR1 that was verified through the *in vivo* binding to AtPIF7 and overexpression in *Arabidopsis*. Most importantly, *DcPAR1* antisense carrot plants showed an impaired taproot development, plastid differentiation and a dramatic decrease in both carotenoids levels and in the relative expression of key carotenogenic genes. These led us to propose that DcPAR1 is a transcriptional cofactor with a key role in the regulation of taproot development and carotenoid synthesis in the carrot taproot grown underground.

## Results

### Carrot plants overexpressing *AtPAR1* present higher carotenoid levels and an increment in *DcPSY1* expression in the taproot grown underground

As a first approximation in order to determine if a functional PAR1 has an effect in carotenoid synthesis in the carrot taproot, we overexpressed *AtPAR1* in carrots. During carrot transformation it trapped our attention that *AtPAR1* embryos presented an orange phenotype instead of the normal whitish coloration in WT embryos (Figure 1A). This early observation suggested that *AtPAR1* could promote the synthesis of carotenoids in carrots in early stages of development. Next, four adult plants (grown 4 months in the greenhouse), that overexpressed *AtPAR1*, were selected, established as independent transgenic lines and subjected to molecular and biochemical analysis (Figure 1B and 1C). Among the four lines, two of them presented higher levels of transgene relative expression in the taproot (OE1 and OE2, Figure 1B), which also showed an increment in *DcPSY1* expression relative to WT taproots (Figure 1B). On the contrary, *DcPSY2* showed a decrease in the relative expression (Figure 1B) whereas the endogenous *DcPAR1* was expressed at similar or reduced levels than WT plants (Supplemental Figure 1). Importantly, transgenic OE1 and OE2 lines, which have the highest expression levels of *AtPAR1* and *DcPSY1*, presented an average of 2.5 times more total carotenoids in their taproots compared to WT plants (919.6 μg/g DW and 796.1 μg/g DW of carotenoids, respectively) (Figure 1C). Moreover, both OE1 and OE2 lines present also higher β-carotene level (2.3 and 2.4 times more than WT roots) and only the OE1 line has significantly higher levels of α-carotene in their taproots (Figure 1C). These results suggest that *AtPAR1* is positively regulating carotenoid synthesis by promoting the expression of the *DcPSY1*, but not *DcPSY2*.

**Figure 1.**
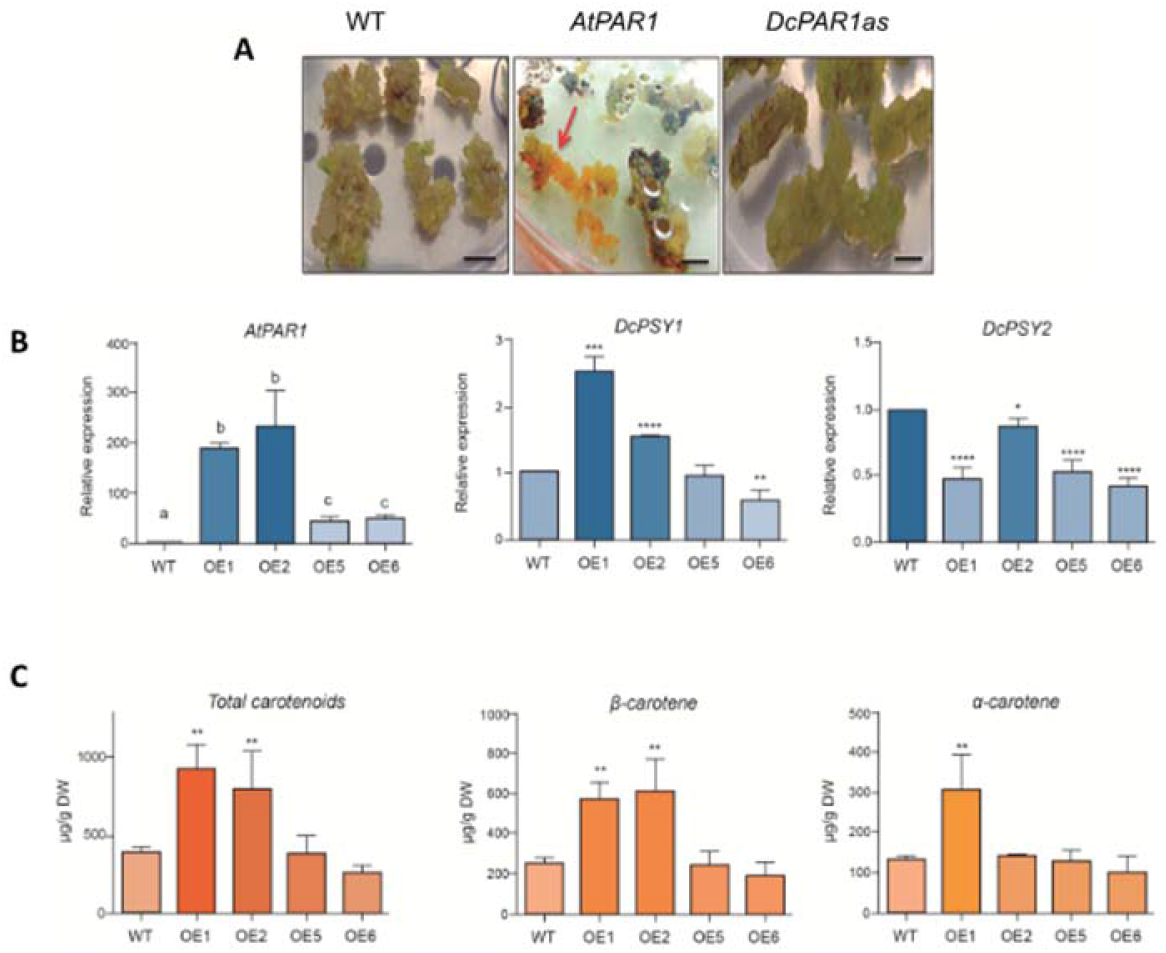
Molecular and biochemical analysis of *AtPAR1* transgenic carrots. **(A)** *In vitro* regeneration of carrot transformant plants. Embryo regeneration stage is shown (18-20 weeks after explants transformation). The arrow indicates orange embryos that later gives rise to whole carrot plants in wild-type (WT), AtPAR1 (AtPAR1:GFP) and DcPAR1as. Scale bar: 5 mm. **(B)** Real time expression of *AtPAR1*, *DcPSY1* and *DcPSY2* in transgenic (OE1, OE2, OE5 and OE6) and wild-type (WT) carrot taproots. Ubiquitin was used as housekeeping gene. **(C)** Total carotenoids, β-carotene and α-carotene levels in transgenic (OE1, OE2, OE5 and OE6) and WT carrot taproots. Letters and asterisks indicate significant differences between WT and transgenic lines (one-way ANOVA analysis with Dunnet post-test; **** p < 0.0001, *** p < 0.001, **p <0.01, * p<0.1)

### DcPAR1 interacts with AtPIF7 in the nucleus

The effect of *AtPAR1* overexpression on taproot *DcPSY1* expression and carotenoid levels suggests that the product of a carrot *PAR1* homologue might have the same role as AtPAR1 in *A. thaliana*. Therefore, we identified contig 42760 (292 bp) obtained from the carrot transcriptome (Arias et al., 2020) which presented a 98% identity with *AtPAR1* (At2g42870, Access N° NM_129848.3) and obtained the complete CDS sequence of *DcPAR1* (Access N° XM_017390696.1) by searching in NCBI (Supplemental Figure 2). The complete CDS of *DcPAR1* (363 bp) presents a 57.7% nucleotide identity with *AtPAR1* and 44% identity at the amino acid level (Figure 2A). We kept more attention to the bHLH domain considering that the AtPAR1 protein presents a bHLH domain that allows it to interact with PIFs and with itself but not to DNA (Galstyan et al., 2011, 2012). A conserved domain search analysis was performed predicting a complete bHLH sequence between residues 29 and 107 (Figure 2A). Comparing this predicted sequence with AtPAR1 bHLH, DcPAR1 presented the key conserved Leu23 residue in Helix 1 (Figure 2B) and the conserved hydrophobic residues such as Val, Ala, Leu and Ile in Helix 2, all of them important for dimer formation. In the basic domain it presents an Asp, Glu and Lys at position 5, 9 and 13, respectively, that, similar to AtPAR1 (Figure 2B), also differ from those (His5-Glu9-Arg13) required for DNA binding (Heim et al., 2003). This sequence analysis suggests that DcPAR1 could be able to dimerize with other bHLH proteins but unable to bind DNA, functionally acting as a transcriptional cofactor, as described for AtPAR1 (Galstyan et al., 2011).

**Figure 2.**
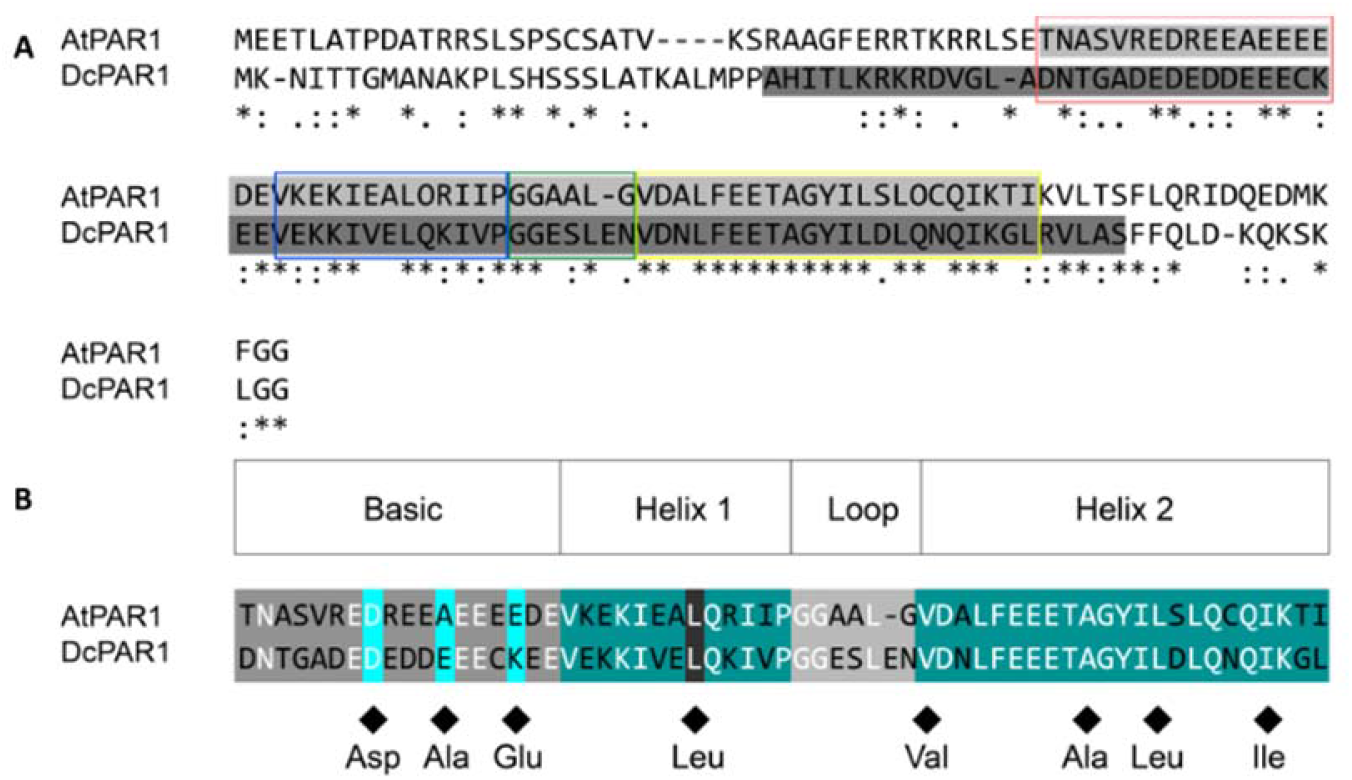
DcPAR1 and AtPAR1 protein alignment. **(A)** The predicted bHLH domains are highlighted in two shades of gray. The red box delimits the basic domains, and blue, green and yellow boxes mark the helix 1, the loop and the helix 2 subdomains, respectively. **(B)** Alignment of AtPAR1 and DcPAR1 bHLH domains. The upper boxes indicate the subdomain extensions based in Heim et al. (2003) and Roig-Villanova et al. (2007). Diamonds highlight residues 5, 9 and 13 (Asp, Glu and Lys, respectively), the conserved leucine residue at position 23 in helix 1 and additional conserved residues in helix 2. Different and identical amino acids are highlighted in each subdomain. All alignments were performed with Clustal Omega.

AtPIF7 has been shown to bind to PHYB (Leivar et al., 2008) and to have a role in promoting Arabidopsis hypocotyls as a result of shade and warm temperature treatments (Li et al., 2012; Fiorucci et al., 2020). Considering that in *A. thaliana* a functional *AtPAR1* binds *in vivo* to PIFs and other bHLH proteins in the nucleus (Hao et al., 2012; Cifuentes-Esquivel et al., 2013; Paulišić et al., 2021), we aimed to determine whether the DcPAR1 protein is nuclear and capable of heterodimerizing with other bHLH proteins. First, we observed that DcPAR1-GFP presents nuclear localization in cells of *Nicotiana benthamiana* leaves (Supplemental Figure 3). Next, we tested interaction of DcPAR1 to AtPIF7 *in vivo* in *N. benthamiana* cells by bimolecular fluorescence complementation (BiFC). In Figure 3 the reconstitution of YFP fluorescence is observed into the cell nucleus of leaves co-agroinfiltrated with the N-terminal part of the YELLOW FLUORESCENCE PROTEIN (YFP, NY) fused to AtPIF7 (NY-AtPIF7) and the C-terminal part of the YFP (CY) fused to DcPAR1 (CY-DcPAR1), thus demonstrating that both proteins interact *in vivo*. Fluorescence is also detected in NY-DcPAR1 and CY-DcPAR1 co-agroinfiltrations (Figure 3), indicating that DcPAR1 can also homodimerize as has been described for AtPAR1 (Bou-Torrent et al., 2015). Together, these results suggested that DcPAR1 has the ability to dimerize with itself and with PIFs, as it has been shown for AtPAR1.

**Figure 3.**
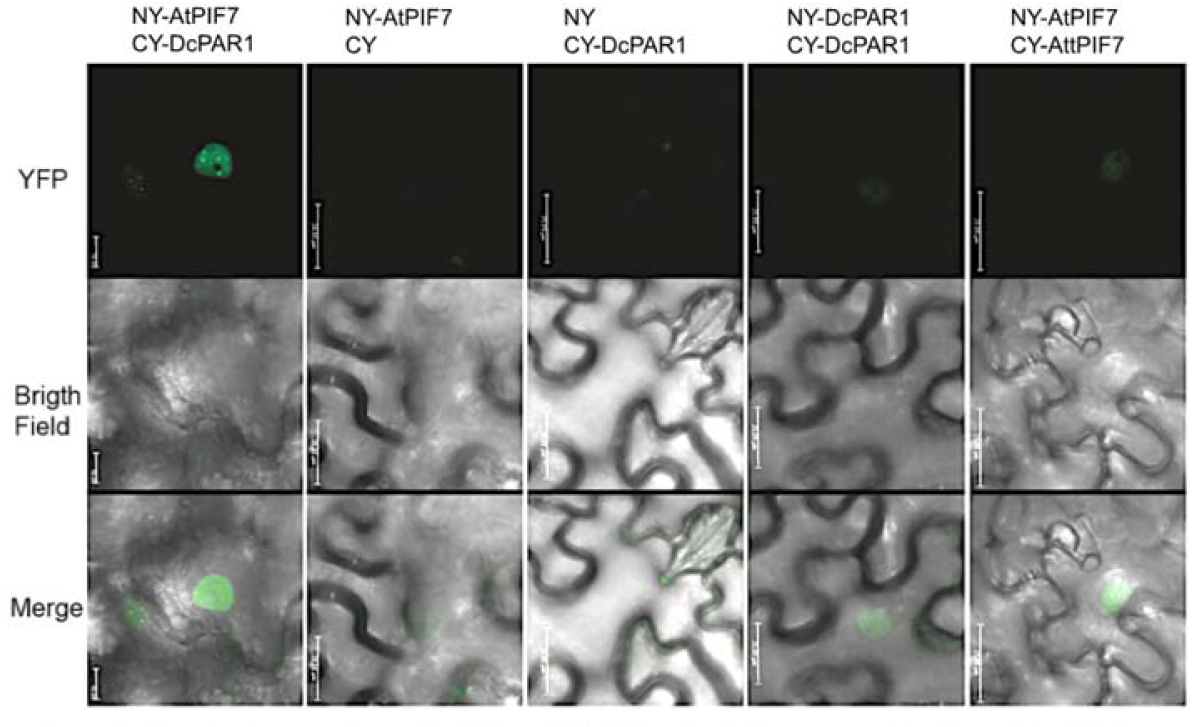
Interaction *in vivo* between DcPAR1 and AtPIF7. Bimolecular Fluorescence Complementation (BiFC) assay between DcPAR1 and AtPIF7 is shown. The NY-AtPIF7/DcPAR1 corresponds to *AtPIF7* or *DcPAR1* genes cloned in fusion to N-terminal region of YFP gene (NY) and CY-AtPIF7/DcPAR1 corresponds to the same genes cloned in fusion to the C-terminal region of YFP gene (CY). The NY and CY are empty vectors. *Nicotiana benthamiana* (tobacco) leaves were co-infiltrated with the NY-and CY-derived plasmids to express the indicated fusion proteins. The top row shows the fluorescence of reconstituted YFP (indicative of a positive interaction of YN and YC fusions), the middle row shows the bright filed of the same area and the bottom row shows the merge of previous conditions. Scale bar: 50 μm

### Arabidopsis plants that express *DcPAR1* present an increment in carotenoids with a higher *AtPSY* expression and AtPSY protein abundance

Overexpression of *AtPAR1* results in dwarf plants and increments in *AtPSY* expression and carotenoid level in Arabidopsis (Roig-Villanova et al., 2007; Bou-Torrent et al., 2015). We next overexpressed *DcPAR1* in Arabidopsis to evaluate if it provides similar phenotypes as *AtPAR1*. We obtained more than 20 independent transgenic lines and selected four T3 transgenic lines that express *DcPAR1* (Figure 4A) for further analysis that were done in two-week-old plants grown under W (long-day photoperiod). The relative expression of the endogenous *AtPAR1* was not significantly affected in the transgenic lines except for L1, which presented a higher expression level than the wild-type line (Col-0) (Figure 4A). Importantly, all transgenic lines presented a significant increase in *AtPSY* relative expression (Figure 4A), as in *AtPAR1* overexpressing lines (Bou-Torrent et al., 2015). The increase in the relative expression of *AtPSY* was accompanied by an increase in the abundance of PSY protein in all transgenic lines (Figure 4B, Supplemental Figure 4 shows coomassie blue staining as control) and an increment in total carotenoids compared to Col-0 (Figure 4C) suggesting that *DcPAR1* positively regulates the expression of *AtPSY* and thus increasing carotenoids content in Arabidopsis.

**Figure 4:**
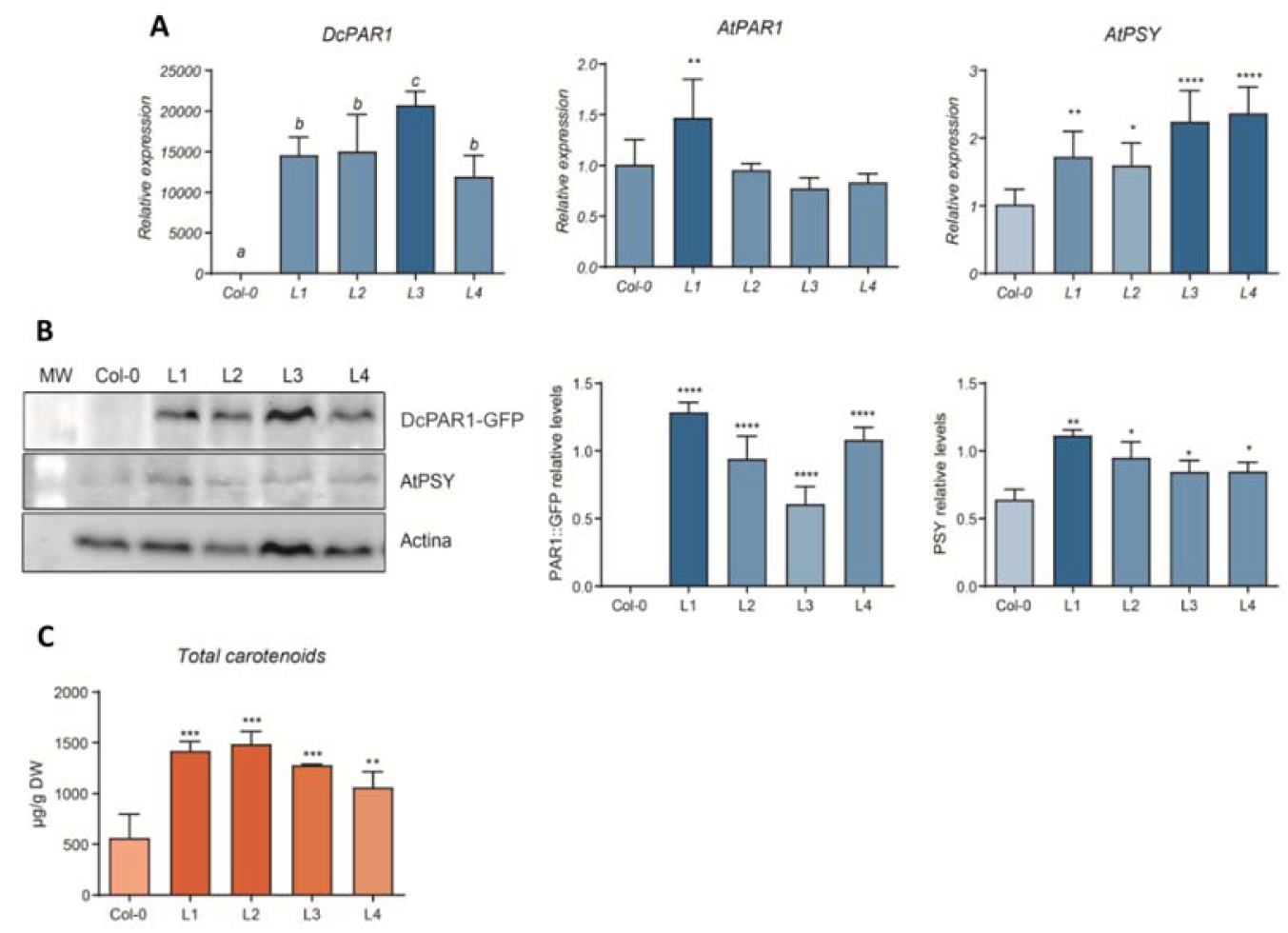
Expression and carotenoid quantification of transgenic Arabidopsis lines overexpressing DcPAR1. **(A)** Relative expression level of *DcPAR1*, *AtPAR1* and *AtPSY* in two-week-old 35S:DcPAR1-GFP (L1, L2, L3, L4) transgenic lines. The relative expression levels were normalized to housekeeping *EFf1A4*. **(B)** Western blot of DcPAR1-GFP, AtPSY and Actin in the transgenic Arabidopsis lines indicated in **A** using antiGFP, antiPSY and anti-ACTIN antibodies. Graphs present the semi-quantification of DcPAR1-GFP and AtPSY from Western blot using Actin abundance as normalizer. Each graph shows the average of three biological replicates. ImageJ program was used for WB semi quantification. MW: molecular weight; Col-0: wild type plants; L1-L4: transgenic plants. **(C)** Total carotenoid content in two-weeks-old Col-0 and transgenic plants. Concentration is presented in μg/g of dry weight (DW). Asterisks indicate significant differences between WT and transgenic plants performed by one-way ANOVA analysis with Dunnet post-test; **** p < 0.0001, *** p < 0.001, **p <0.01, * p<0.1.

To determine if DcPAR1 also affects elongation, we assessed phenotypic analysis in two-week- and 1.5-month-old transgenic lines. In two-week-old seedlings, transgenic lines presented a reduced length in hypocotyl, cotyledons and primary leaves length (Supplemental Figure 5) similar to that reported for Arabidopsis *AtPAR1* overexpressing lines (Roig-Villanova et al., 2007; Hao et al., 2012; Zhou et al., 2014; Bou-Torrent et al., 2015). Moreover, all transgenic lines present a dwarf phenotype (Figure 5A) with a significant reduced stem height and silique length (Figure 5B) at the mature stage. These results suggest that the *DcPAR1* is functional *in vivo* in promoting carotenoid synthesis and inhibiting growth in *A. thaliana*.

**Figure 5:**
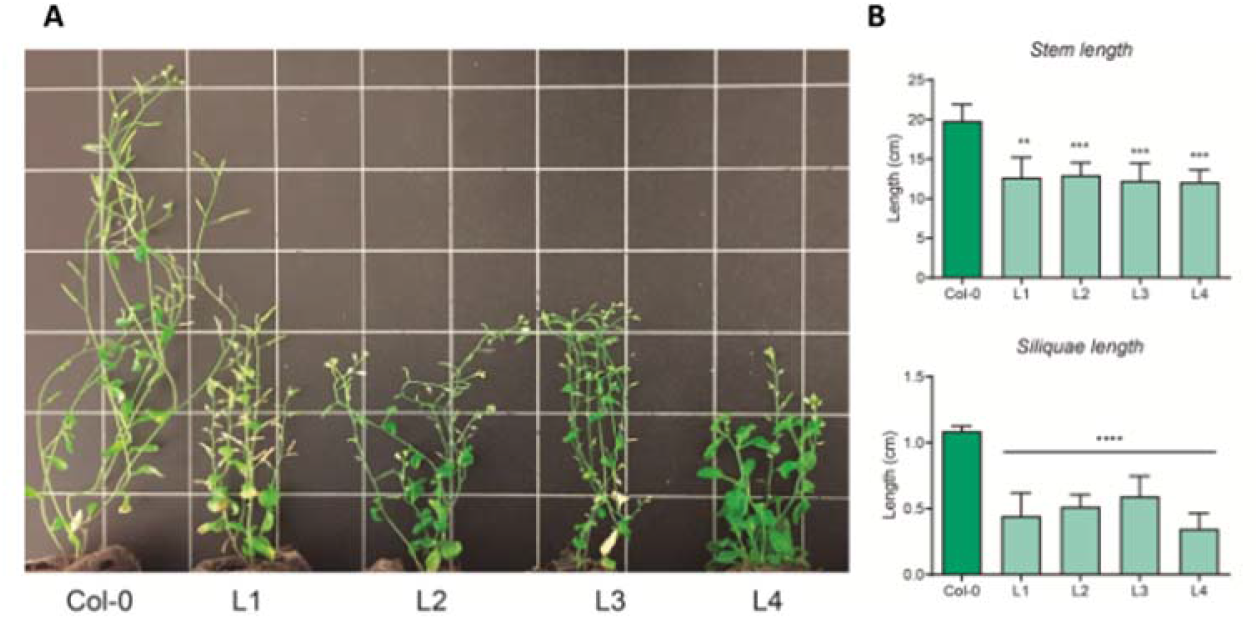
Phenotype of mature transgenic Arabidopsis lines overexpressing DcPAR1. **(A)** Picture showing the length and phenotype of representative six-weeks-old plants of Col-0 and T3 transgenic lines. **(B)** Stem and **(C)** Silique length of six-week-old Col-0 and T3 transgenic plants. Each value is the mean ± SE/SD of 5 plants each and using ImageJ program. Asterisks indicate significant differences between WT and transgenic plants performed by one-way ANOVA analysis with Dunnet post-test; **** p < 0.0001, *** p < 0.001, **p <0.01, * p<0.1.

### Repression of *DcPAR1* expression affects carrot taproots growth, carotenoid composition and carotenogenic gene expression

The next step was to evaluate the role of *DcPAR1* in carrot. For this, transgenic antisense carrot plants for *DcPAR1* were generated. During *in vitro* regeneration process of DcPAR1 antisense plants *(DcPAR1as)* no phenotypic differences were observed respect to WT embryogenesis (Figure 1A). Eleven mature plants were obtained, and the molecular analysis confirmed that ten of them were transgenic (Supplemental Figure 6). Four transgenic lines were selected for further molecular and biochemical analysis. As is shown in Figure 6A, all transgenic lines presented a dramatic reduction between 80% and 99% in the *DcPAR1* expression and a drastic reduction in the taproot secondary growth at the mature stage, as shown for line AS10 (Figure 6B). The taproots of a four-month-old representative line that grew underground was considerably smaller and thinner than WT taproots of the same age and grown under the same conditions. The average mass of the transgenic roots did not exceed 400 mg while the average mass of a four-month-old WT carrot taproot can weigh up to 7 g (Figure 6B). In addition, the roots were pale suggesting that a significant reduction in carotenoid levels might have occurred (Figure 6B). In addition, expression of key genes required for carotenoid synthesis, *DcPSY1*, *DcPSY2*, *DcLCYE*, *DcLCYB2*, *DcCHXB1* and *DcCHXB2*, was significantly lower in all silenced lines relative to the WT, showing a reduction between 40% and 99% (Figure 6C). Moreover, a 50%-80% decrease in total carotenoid levels in all silenced lines was also obtained (Figure 7A). Likewise, all the transgenic lines showed a 4- to 60-fold decrease in α-carotene and 4- to 58-fold in β-carotene (Figure 7B). However, an unexpected 6- to 15-fold increase in lutein levels was obtained (Figure 7B). This pigment does not normally accumulate in the orange carrot taproot that grows underground but in photosynthetic tissues and in the WT carrot roots that grow exposed to light (Fuentes et al., 2012).

**Figure 6:**
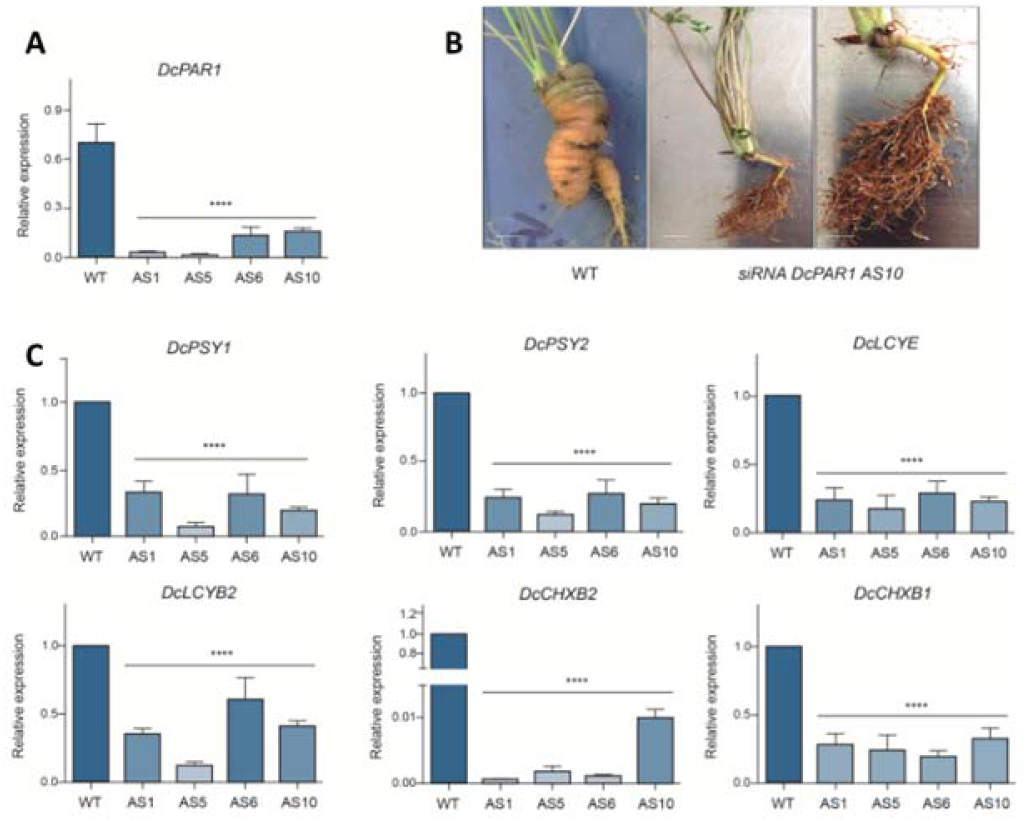
Phenotype and expression analysis of four-months-old DcPAR1as transgenic carrot taproots. **(A)** Relative expression level of *DcPAR1* in AS1, AS5, AS6 and AS10 transgenics lines. **(B)** Representative phenotype of a WT and a DcPAR1as transgenic carrot root (line AS10) grown four months underground after transplanting. Bar: 1 cm. **(C)** Relative expression level of *DcPSY1*, *DcPSY2*, *DcLCYB2*, *DcLCYE DcCHXB1* and *DcCHXB2*. The relative expression was carried out in triplicate (two technical replicas each) and normalized to the housekeeping *Ubiquitin* gene. WT was used as calibrator. Asterisks indicate significant differences between transgenic lines and WT plant determined by one-way ANOVA analysis with Dunnet post-test; **** p < 0.0001, *** p < 0.001, **p <0.01, * p<0.1.

**Figure 7:**
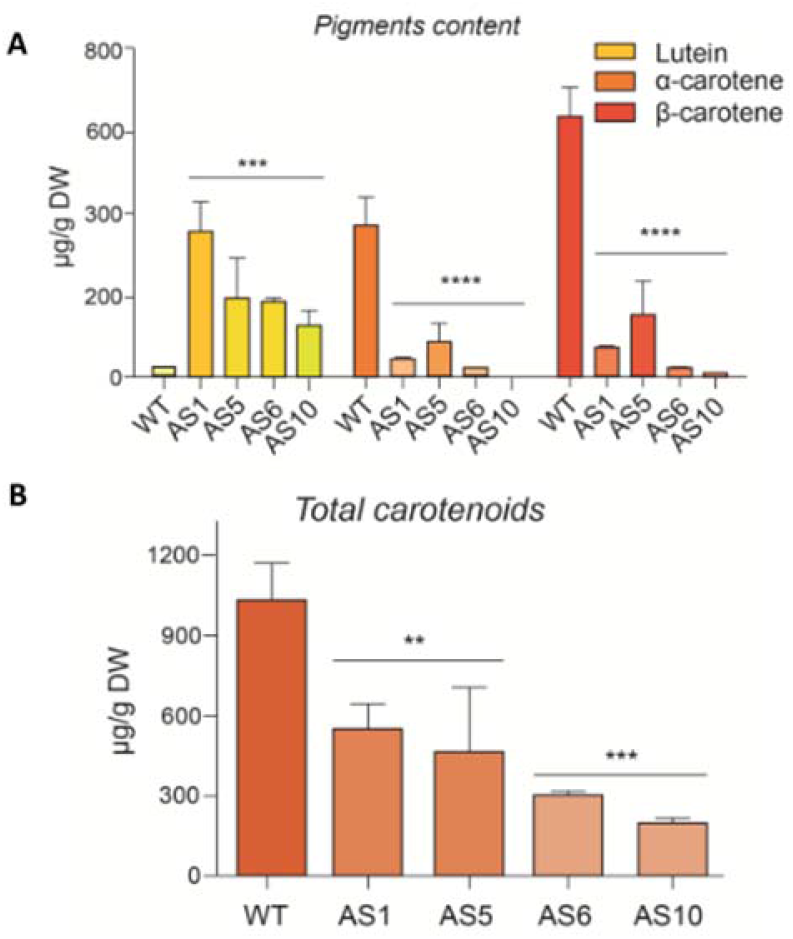
Total and individual carotenoid content in four-month-old DcPAR1as carrot taproots. **(A)** lutein, α-carotene and β-carotene levels was determined by HPLC-RP and **(B)** total carotenoids were quantified in a spectrophotometer at 474 nm. The pigment content is indicated in μg/g of dry weight (DW) and was measured in triplicate. Asterisks indicate significant differences between transgenic lines and WT plant determined by one-way ANOVA analysis with Dunnet post-test; **** p < 0.0001, *** p < 0.001, **p <0.01, * p<0.1

With the purpose of verifying that the taproot phenotype in silenced plants persisted in time, the plants were grown four additional months underground in the greenhouse (except the AS10 line that could not be maintained because it was harvested and could not be recovered). We observed that, although the size of transgenic roots increased respect to the four months, they were still smaller and thinner than the WT taproots and presented a greenish coloration in the external part of the taproot and a yellowish coloration in the taproot centre (Figure 8A). The relative expression level of *DcPAR1* in 8-months transgenic lines kept reduced between 60% to 75% (Figure 8A). Likewise, the relative expression of the carotenogenic genes in transgenic lines were still lower than in WT plants (Figure 8B), confirming that *DcPAR1* indirectly regulates the expression of carotenogenic genes in carrot root.

**Figure 8:**
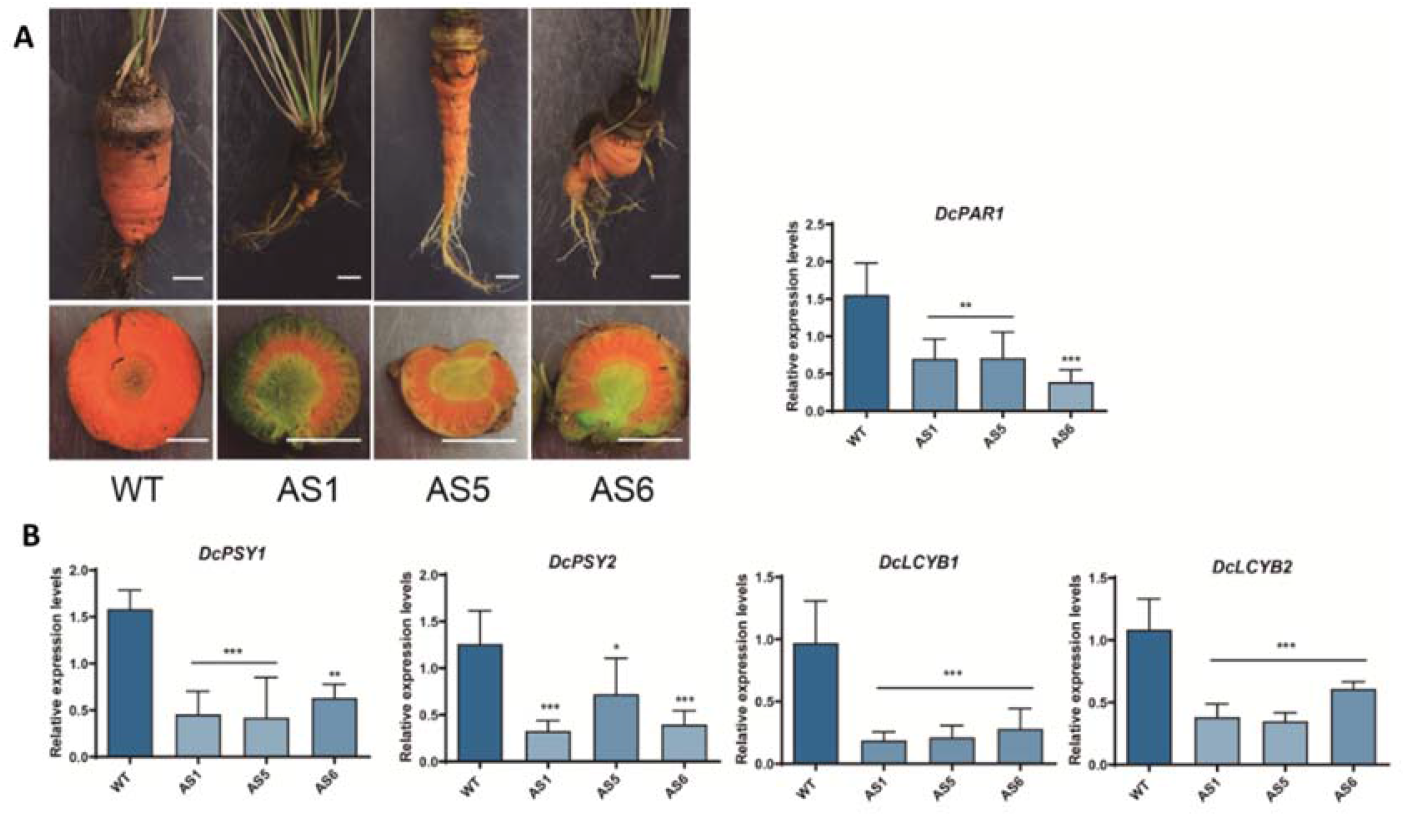
Phenotype and expression analysis of eights-months-old DcPAR1as transgenic carrot taproots. **(A)** Representative phenotype of carrot taproots of a WT and AS1, AS5 and AS6 DcPAR1as transgenic plants grown eight months underground after transplanting (line AS10 was harvested at four months and not shown in here). Bar: 1 cm. Graph in the right shows the relative expression level of *DcPAR1* in the indicated lines. **(B)** Relative expression level of *DcPSY1*, *DcPSY2*, *DcLCYB2*, *DcLCYE*, *DcCHXB1* and *DcCHXB2*. Relative expression was carried out in triplicate (two technical replicas each) and normalized to the housekeeping *Ubiquitin* gene. Asterisks indicate significant differences between transgenic lines and WT plant determined by one-way ANOVA analysis with Dunnet post-test; **** p < 0.0001, *** p < 0.001, **p <0.01, * p<0.1

Carotenoids and chlorophylls quantification showed that DcPAR1as lines presented 2.4-fold fewer total carotenoids (Figure 9A) and an increment of 3-5 to 8-fold in chlorophyll a and b reaching 100-250 μg/g DW (Figure 9B), suggesting that the absence of PAR1 leads to chloroplast instead of chromoplasts differentiation. Likewise, transgenic antisense plants present a 3.5 to 14-fold reduction in α-carotene and 3 to 9-fold reduction in β-carotene levels respect to WT plants. As in younger taproots, a significant increment in lutein was also observed, which reached up to 200-250 μg/g DW (compared to 30 μg/g DW in WT) (Figure 9C), thus explaining the yellowish coloration of antisense transgenic taproots respect to the orange pigmentation of WT (Figure 8A). Taking all the results together, we suggest that *DcPAR1* is a functional cofactor that has an important regulatory role not only in carotenoid synthesis in carrot taproot that grows underground, but also in secondary taproot development and plastid differentiation.

**Figure 9:** Carotenoids and chlorophylls content in eight-months-old DcPAR1as transgenic carrot taproots. **(A)** Total carotenoids levels and **(B)** cholorophyll a and b in carrot taproots of WT and AS1, AS5 and AS6 DcPAR1as transgenic plants. **(C)** Lutein, α-carotene and β-carotene levels was determined by HPLC-RP. Pigment content is indicated in μg/g of dry weight (DW) and was measured in triplicate. Asterisks indicate significant differences between transgenic lines and WT plant determined by one-way ANOVA analysis with Sidak post-test; **** p < 0.0001, *** p < 0.001, **p <0.01, * p<0.1

## Discussion

### *DcPAR1* boosts carotenoid synthesis and promotes a photomorphogenic phenotype in Arabidopsis

Carotenoid biosynthesis regulation in plants has been extensively studied. Although there is much information about how this process is regulated by several factors, there is still much to explore especially respect to how light regulates this process in plant organs that normally grow with no direct W, such as roots (Stanley and Yuan, 2019; Quian-Ulloa and Stange, 2021). With the purpose to understand how carotenoids are synthesized in orange carrot taproot, we compared through RNAseq the expression profile of carrot roots that grew in W respect to those that grew underground (Arias et al., 2020). We identified several photomorphogenic genes that were upregulated in the underground condition and among them *DcPAR1*, *DcPHYA* and *DcPIF4* presented twice as much expression level in darkness as in the root that grew in light (Arias et al., 2020). PAR1 is a recognized negative regulator factor in the shade avoidance syndrome (SAS) and a positive factors during seedling de-etiolation in Arabidopsis (Roig-Villanova et al., 2007; Hao et al., 2012; Zhou et al., 2014). In shade, the expression of *PAR1*, *PAR2* and the *LONG HYPOCOTYL IN FAR RED 1* (*HFR1*) is induced, factors that encode transcriptional bHLH cofactors that participate in the compensation of excessive hypocotyl elongation during SAS (Roig-Villanova et al., 2007; Bou-Torrent et al., 2008; Hao et al., 2012; Zhou et al., 2014). From these factors, PAR1 interacts with several bHLH proteins that act as positive regulators of SAS such as BEEs, BIMs and PIFs by partially preventing the binding of these transcription factors to LREs of photomorphogenic genes (Roig-Villanova et al., 2006; Cifuentes-Esquivel et al., 2013; Ballaré and Pierik, 2017; Roig-Villanova et al., 2007; Hao et al., 2012; Stange and Flores, 2012; Zhou et al., 2014; Bou-Torrent et al., 2015; Fernández-Milmanda and Ballaré, 2021; Quian-Ulloa and Stange, 2021). Importantly, PAR1 promotes *PSY* expression under shade (Bou-Torrent et al., 2015) and, of the positive SAS regulators BEEs, BIMs and PIFs, only PIFs participate in repressing *PSY* expression and carotenoid synthesis in the shade (Toledo-Ortiz et al., 2010). PIFs remain stable in the shade condition and bind to the LRE motifs (Lorrain et al., 2008; Hornitschek et al., 2012).

AtPAR1 is an nuclear and atypical bHLH, presents a functional HLH protein domain, but it lacks a functional DNA-binding domain, so its regulatory function is performed through the dimerization with PIFs transcription factors. To date the direct interaction between AtPAR1 and AtPIF4 and AtPIF1 has been shown (Roig-Villanova et al., 2007; Hao et al., 2012; Bou-Torrent et al., 2015).

Our bioinformatic analysis indicated that DcPAR1 has a similar structural pattern to AtPAR1, especially at the bHLH domain, suggesting that it could fulfill similar functions. Importantly, the key amino acids at the basic domain that determines the binding to DNA are also replaced by similar amino acids that are present in AtPAR1 (Heim et al., 2003, Figure 2B). Likewise, the essential amino acid that determine the protein-protein interaction are still conserved in DcPAR1 (Heim et al., 2003, Figure 2B) which in functionality terms allows DcPAR1 to interact with other bHLH proteins, such as itself and AtPIF7 (Figure 3), an essential characteristic for a functional PAR1.

AtPAR1 interacts with AtPIF1, and consequently induces the expression of *AtPSY*, thus increasing the content of carotenoids together with the promotion of a photomorphogenic phenotype (Roig-Villanova et al., 2007; Bou-Torrent et al., 2015). Interestingly, Arabidopsis plants overexpressing *DcPAR1-GFP* and grown in W, presented a stable accumulation of DcPAR1-GFP, suggesting that it is stable in W, similarly to AtPAR1 that is stable in low and high R:FR, but not in the dark (Zhou et al., 2014). The higher DcPAR1-GFP accumulation correlates with an increase in *AtPSY* relative expression levels and in PSY protein with a final rise in total carotenoids (Figure 4C), which strongly suggest a positive role of DcPAR1 on carotenoids synthesis in Arabidopsis. Indeed, the expression of the endogenous AtPAR1 was not affected suggesting that PAR1 does not induce its own expression in Arabidopsis. It remains to be elucidated whether the DcPAR1 protein would be promoting *AtPSY* transcription through binding exclusively to PIF-type transcription factors or if there are other transcription factors that may be involved in this process, such as BEEs and BIMs (Cifuentes-Esquivel et al., 2013).

Moreover, according to the dwarf phenotype of Arabidopsis lines overexpressing *AtPAR1* (Roig-Villanova et al., 2007), 2 weeks-old *DcPAR1* transgenic lines presented several characteristics of a pronounced photomorphogenic phenotype (Supplementary Figure 5) that remains until 1.5 months (Figure 5). Given these results we strongly suggest that *DcPAR1* and *AtPAR1* are orthologs.

### *DcPAR1* positively regulates carotenoid synthesis and plastid differentiation in carrot taproot

Knowing that *AtPAR1* positively regulates the carotenoids synthesis in Arabidopsis photosynthetic tissue (Roig-Villanova et al., 2007; Bou-Torrent et al., 2015) it seemed a good starting strategy to visualize the *in vivo* role of a functional PAR1 in carotenoids synthesis in carrot taproot.

Carrot *AtPAR1-GFP* lines with the highest transgene expression (OE1 and OE2) presented a significant increment in the relative expression levels of *DcPSY1*, but not in *DcPSY2* and *DcPAR1*, in correlation to an increase in carotenoid levels in the taproot of mature carrots (Figure 1), suggesting that AtPAR1:GFP is sufficient to produce the carotenoid increment in these lines. Interestingly, the transgenic lines with the lower transgene expression level (OE5 and OE6), produced a significant reduction in *DcPSY1* (except for OE5), *DcPSY2* and *DcPAR1* expression but without affecting the level of carotenoids, which remains similar to the WT. The decrease in the relative expression of *DcPAR1* may be an endogenous compensation mechanism leading to speculate that AtPAR1 may regulate the expression of its ortholog gene in carrot. Given these results, we suggest a positive regulation of AtPAR1 on *DcPSY1* and not on *DcPSY2* and that the effect generated is due to the expression of *AtPAR1* and not of the endogenous gene.

The regulation of AtPAR1 on the different *DcPSY* paralogs may depend on the transcription factors that bind to their promoters that can be regulated by AtPAR1. It could be possible that AtPAR1 dimerizes to some endogenous carrot PIFs that bind preferably to *DcPSY1* than to *DcPSY2* promoter, and the higher AtPAR1 abundance may recruit this type of PIFs preventing their binding to *DcPSY1* promoter. Indeed, light-responsive elements (LRE) are located in the *DcPSY1* and *DcPSY2* promoters (Simpson et al., 2018). It may be interesting to determine which transcription factors directly bind to the promoters of *DcPSY1* and *DcPSY2* in carrots. Since *DcPSY2* is mostly induced by salt stress and ABA and that AREB transcription factors are able to bind to *DcPSY2* promoter (Simpson et al., 2018), we suggest that *DcPSY2*, and not *DcPSY1*, is induced under abiotic stress (Simpson et al., 2018). It remains to be determined whether *DcPSY1* could be most associated to carotenoid synthesis during taproot development. On the other hand, in transgenic carrots that express *AtDXS* (a gene that participates in the synthesis of the metabolic precursors for carotenoids) an increase in carotenoid levels was observed also in correlation to an increase in the expression of *DcPSY1* and *DcPSY2* (Simpson et al., 2016b) suggesting that both genes determine the levels of carotenoids in carrot taproots.

Likewise, the post-transcriptional gene silencing of *DcPAR1* in carrots is a better strategy to determine the role of the endogenous gene in carrots. *DcPAR1* antisense had important consequences on the carotenoid levels (Figure 7 and 9), which decrease significantly in correlation to a decrease in the expression of all carotenogenic genes analyzed, including both, *DcPSY1* and *DcPSY2* (Figure 6 and 8). Similar results were reported in *DcLCYB1* antisense carrots (Moreno et al., 2013), where a decrease of 40-80% in total carotenoid levels together with a decrease in the expression levels of *DcPSY1* and *DcPSY2* was observed (Moreno et al., 2013). Moreover, *DcPAR1* antisense lines showed a dramatic reduction in the expression on all carotenogenic genes evaluated, in a direct or indirect manner. Possibly, DcPAR1 has the ability to bind to PIFs or other unknown transcription factors some of which would directly down or upregulate *DcPSY1*, *DcPSY2*, *DcLCYB1*, *DcLCYE*, *DcCBHX1* and/or *DcCBHX2*. It is also possible that DcPAR1-DcPIFs complex exerts a direct regulation upon *DcPSY*s and the activation of them has a direct impact on the other genes in the pathway. To establish a more in-depth explanation in this regard, it is necessary to determine which other factors, in addition to PIFs, are binding to PAR1 in carrots during development and which transcription factors bind to promoters of carotenogenic genes, inducing their expression. In Arabidopsis, ELONGATED HYPOCOTYL 5 (HY5) binds to PIFs and also competes with PIFs for binding to LRE, promoting *PSY* expression and inducing the differentiation of etioplasts into chloroplasts with an increment in carotenoids and chlorophylls in plants (Von Lintig et al., 1997; Ronen et al., 1999; Woitsch and Römer, 2003; Toledo-Ortiz et al., 2010). It has been proposed that there would be a genetic interaction between AtPAR1 and AtHY5 because double antisense-PARs/*hy5* mutants present a more elongated hypocotyl than simple mutants (Zhou et al., 2014). Therefore, HY5 role in PAR1 antisense lines could be explored. Nevertheless, these results support the hypothesis that DcPAR1 would be a positive regulator of carotenoid synthesis in carrot taproot.

Carrot DcPAR1as lines produce a significant increment in lutein (in four- and eight-months old carrot taproots) similar to WT carrot roots grown in light, but absent in taproots grown underground (Fuentes et al., 2012; Arias et al., 2020). Most confusing is that *DcCHXB1* and *DcCHXB2*, involved in lutein synthesis are deeply down regulated (Figure 6C), suggesting that the enzymes may be stabilized by an unknown mechanism.

In addition, DcPAR1 antisense lines present an increase in chlorophylls (in eight-months old carrot taproots) that are normally absent in WT taproots, suggesting that DcPAR1 not only is required for carotenoid synthesis or accumulation but also in chromoplast differentiation. Indeed, DcPAR1 antisense lines present chlorophylls as the WT roots that were grown in W (Fuentes et al., 2012) in which *DcPAR1* expression is lower than in the taproot grown underground (Arias et al., 2020).

### *DcPAR1* positively regulates carrot taproot development

It has been shown that the accumulation of carotenoids in carrot roots is genetically associated to a homologue of the Arabidopsis *PSEUDO-ETIOLATION IN LIGHT* (*PEL*) gene (Iorizzo et al., 2016). PEL presumably acts as a repressor of photomorphogenesis. Carrot varieties with a loss-of-function allele of the *PEL* gene accumulate carotenoids in the root, suggesting that high pigment contents might result from a derepressed development of carotenoid-accumulating plastids (i.e., chloroplasts in the light but chromoplasts in the dark) (Llorente et al., 2017). Our results support this possibility and go further in proposing DcPAR1 as an antagonistic actor of PEL1. Indeed, the expression of *AtPAR1* in carrots produces orange embryo showing an early and positive effect of PAR1 on carotenoid synthesis (Figure 1) but it did not generate an evident change in photomorphogenic phenotype at the mature stage.

Although *AtPAR1* antisense produced elongated and hyper-etiolated plants (Zhou et al., 2014), carrot DcPAR1 antisense lines presented similar embryogenic and photomorphogenic development that control plants, especially in seedlings hypocotyl length (not shown), although they presented up to 95% of *DcPAR1* reduced expression. However, the consequences in the taproot of *DcPAR1* antisense adult plants were drastic, showing a dramatic reduction in the size (thickness and length) and weight of the taproot in 4-month-old plants, a tendency that was maintained at 8 months of development. The phenotype obtained is partly reminiscent of that observed when silencing *DcLCYB1*, where the roots generated were thinner than the controls (Moreno et al., 2013) but less drastic than DcPAR1 antisense, showing a relevant role for DcPAR1 in carrot taproot development.

Carrot root development has been previously reported to be closely related to carotenoid synthesis (Suslow et al., 1999; Clotault et al., 2008). Thus, it is possible that both processes are being regulated by similar factors at the transcriptional or post-transcriptional level. PAR1 seems to be a good candidate to participate in both processes. These results and those reported in Arabidopsis suggest that PAR1 has an important regulatory role on carotenoid synthesis and development. Considering that PAR1 is upstream of the pigment synthesis pathway, it has a broader functional role on the carotenoid synthesis in parallel with the regulation of root development in the absence of W. AtPAR1 reduces hypocotyl elongation in shade through the repression of auxin-induced genes such as *SAUR15* and *SAUR68* likely through their ability to inhibit DNA binding of PIFs or other bHLH to its binding motifs in the *SAUR15* and *SAUR68* promoters (Roig-Villanova et al., 2006, 2007; Bou-Torrent et al., 2008, 2015; Hao et al., 2012; Zhou et al., 2014). Moreover, some of the phenotypes of AtPAR1 overexpressing lines are similar to those reported in auxin, brassinosteroid or gibberellin mutants (Nakazawa y cols., 2001). Considering that DcPAR1 antisense lines present a short and thin taproot with more lateral root abundance we propose that DcPAR1 may promote auxin metabolism de-repression. Therefore, it remains to know if DcPAR1 could have a role in hormone signaling during taproot development.

Based on our results, we propose a simple model regarding DcPAR1 role in carotenoid synthesis, plastid differentiation and taproot development (Figure 10). Exposure of the orange carrot roots to light (R/L) decreases *DcPAR1* transcripts levels which impairs the expression of carotenogenic genes such as *DcPSYs*. This leads to a reduced amount of PSY protein accumulation, which in turn generated a drop in total carotenoid accumulation with reduced α-carotene and β-carotene levels but an increase in lutein and chlorophylls. This might result in chloroplast (rather than chromoplasts) differentiation and impairment in taproot development. On the contrary, in roots grown underground (e.g., covered with soil; R/D), the higher transcripts levels of *DcPAR1* promotes an increment in *DcPSYs* expression and PSY accumulation that leads to an increment in carotenoids synthesis (specially α-carotene and β-carotene) that would promote chromoplast differentiation and secondary root development. Indeed, recently it was shown that synthetically inducing a burst in the production of phytoene (the product of the PSY enzymes and the first committed intermediate of the carotenoid pathway) elicits an artificial chloroplast-to-chromoplast differentiation in leaves (Llorente et al., 2020). Altogether, these results let us to propose that DcPAR1 is a key factor for secondary root development, plastid differentiation and carotenoid synthesis in carrot taproot grown underground.

**Figure 10:**
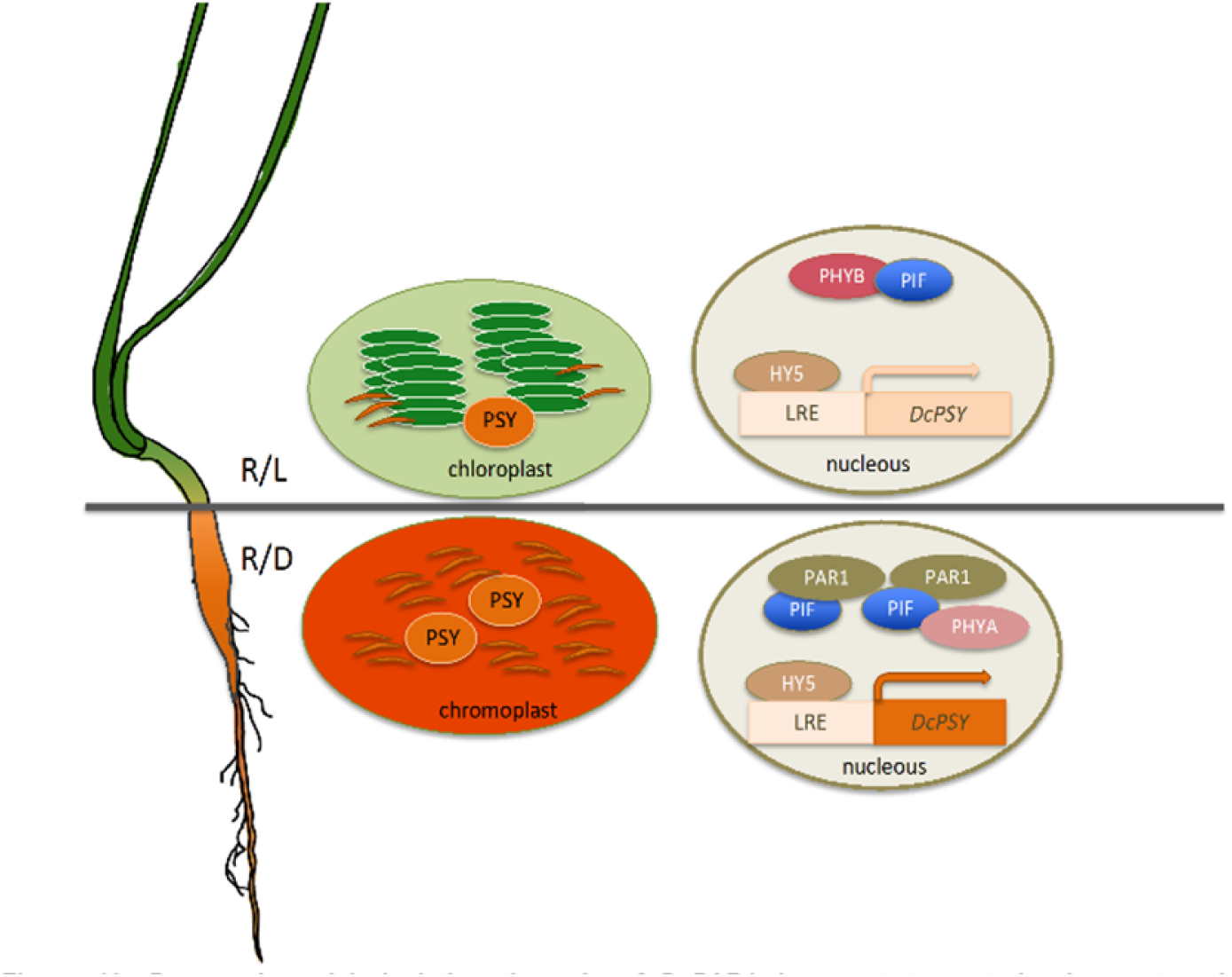
Proposed model depicting the role of DcPAR1 in carrot taproot development and accumulation of carotenoids. Roots grown exposed to W (R/L) presents a reduced expression of *DcPAR1* which causes a decrease in the expression of carotenogenic genes such as *DcPSYs*, and in PSY protein accumulation. As a consequence, total carotenoids, α-carotene and β-carotene accumulation drop but chlorophylls and lutein levels increase according to chloroplast differentiation. Taproot development is also impaired. On the contrary, in roots grown underground (R/D), the enhance transcripts levels of *DcPAR1* promotes an increment in *DcPSYs* expression and PSY accumulation that leads to an increment in carotenoids synthesis, chromoplast differentiation and secondary taproot development.

## Conclusions

- *DcPAR1*, like its Arabidopsis counterpart *AtPAR1*, induces *AtPSY* expression, boosts carotenoid synthesis and produces a dwarf phenotype when overexpressed in *A. thaliana*.
- *DcPAR1* positively regulates carotenoid synthesis and plastid differentiation in carrot taproot grown underground.
- DcPAR1 is a key factor for carrot secondary root development.

## Materials and Methods

### Bioinformatic analysis

NCBI platform was used to obtain the complete CDS of *DcPAR1* using the contig 42760 (202 bp) that corresponds to *DcPAR1* and was obtained from the RNAseq analysis (Arias et al, 2020). Conserved domain search analysis was performed using the InterPro Scan (https://www.ebi.ac.uk/interpro/search/sequence-search) and SMART (http://smart.embl-heidelberg.de/) in order to identify the bHLH functional regions in DcPAR1.

### *DcPAR1* gene amplification and vector construction

The complete CDS sequence of *PAR1* of *Daucus carota* var. Nantaise (363 pb without stop codon) (NCBI access number XM_017390696.1) was amplified from carrot taproot cDNA, using Herculase II Fusion DNA Polymerase (Agilent technologies) and DcPAR1.F and DcPAR1.R primers (Supplementary Table I). The sequence was cloned into the entry PCR^™^8/GW/TOPO^®^ vector (Invitrogen) according to manufacturer’s instructions and sequenced (Macrogen Corp, USA). Clones of PCR8:DcPAR1 in sense and antisense orientation were recombined with the destination vector pGWB5, obtaining the binary vectors pG5_35S:PAR1:GFP (sense DcPAR1-GFP) and pG5_35S:PAR1as (antisense DcPAR1as) that were used to overexpress and silence *DcPAR1*, respectively. The pBF1_35S:AtPAR1-GFP binary vector was selected overexpress *AtPAR1* as a fusion to the *GFP* reporter gene (AtPAR1-GFP) and has been described elsewhere (Roig-Villanova et al., 2007). All binary plasmids were transformed into *Agrobacterium tumefaciens* strain GV3101 for stable Arabidopsis and carrot transformation.

### *Agrobacterium-mediated* transformation of *Daucus carota* and *Arabidopsis thaliana*

Commercial seeds of *Daucus carota* var. Nantaise were surface-sterilized and sown in culture medium MS (4.4 g/L MS salts (Murashige and Skoog, 1962), 20 g/L sucrose and 0,7% agar) and grown in a culture chamber under 16 h long day photoperiod at 23-25 °C. Plants of 10-20 days post-germination were used for *Agrobacterium*-mediated transformation as described in Gonzalez-Calquin and Stange (2020). Briefly, the epicotyls of wild-type plants grown *in vitro*, were cut and co-cultivated in presence of *A. tumefaciens* strain GV3101 carrying the DcPAR1-GFP, DcPAR1as or AtPAR1:GFP vectors and placed on solidified MS in darkness. After two days, the explants were transferred to MS culture medium supplemented with 1 mg/L of 2,4-D, 50 mg/L of Hygromycin (for transgene selection) and 200 mg/L of Timentin (for *Agrobacterium* control) for embryo development. After 4-5 weeks in darkness, the explants were transferred to MS culture medium supplemented with 0.5 mg/L of 2,4-D, 100 mg/L of Hygromycin and 200 mg/L of Timentin and grown in W (16 h long-day photoperiod at 23-25 °C) for 7 weeks approximately. Finally, the hygromycin-resistant embryos were transferred to MS culture medium without hormones to promote elongation of carrot seedlings. Plants of 5-7 cm length were transferred to a mixture of soil: vermiculite (1:1) and maintained under the same photoperiod and temperature conditions described above. Transgenic plants were selected by amplifying the respective transgen form genomic DNA: *AtPAR1* gene in the case of carrot plants transformed with pBF1:AtPAR1:GFP (Bou-Torrent et al., 2015) with AtqPAR1.F and AtqPAR1.R primers (Supplementary Table I), *DcPAR1* gene in the case of Arabidopsis plants using primers DcqPAR1.F and DcqPAR1.R (Supplementary Table I) or including *GFP* amplification in the case of carrot plants transformed with pG5:PAR1as, using primers DcPAR1.R and qEGFP.R (Supplementary Table I).

*A. thaliana* wild-type plants (Col-0 ecotype) were transformed using floral dip (Zhang et al., 2006). Seeds were surface sterilized in a solution of sodium hypochlorite (50% v/v) for 6 min, washed three times with sterile water and sown in solid MS medium (4.4 g/L MS salts, 0.44% vitamins, 0.01% myo-inositol and 0.7% agar pH 5.8 with or without selection). They were kept in a growth chamber with a 16 h long day photoperiod illuminated with cool-white fluorescent light (115 μmol/m^2^s) at 22°C for 2-6 weeks depending on the experimental assay. Selected transgenic lines were transferred to a greenhouse in a mixture of soil:vermiculite (1:1) for molecular analysis. The selection of T3 homozygous transgenic lines was carried out by cultivation of selected transgenic T1, T2 and then T3 seeds in MS medium with hygromycin (15 μg/ml) and selected those T3 with up to 95% survival for further molecular, biochemical and phenotypical analyses.

### Pigments extraction and quantification

In carrot and Arabidopsis, carotenoid and chlorophyll extraction was carried out as described previously (Simpson et al., 2016). Pigments were extracted from 20-100 mg of fresh carrot transgenic taproots and 100 mg of entire 2 weeks-old Arabidopsis T3 lines with 4 ml of hexane:acetone:ethanol (2:1:1 v/v). After centrifugation, the upper phase was recovered and dried with N_2_. For total carotenoids and chlorophylls quantification, pigments were resuspended in 1 mL of acetone and quantified in a spectrophotometer (Shimadzu) at 750, 662, 645 and 474 nm in quartz cuvettes. Absorbance at 662, 645 and 474 nm permitted to determine the concentration of chlorophyll a, chlorophyll b and total carotenoids, respectively. The absorbance at 750 nm determines the turbidity of the sample which may result in underestimation of the pigment concentration. Then, concentration of chlorophyll a, chlorophyll b and total carotenoids was determined as described previously (Lichtenthaler and Buschmann, 2001). Individual carotenoids were quantified in a Shimadzu HPLC (LC-10AT) with a diode array detector using a RP-18 Lichrocart 125-4 reverse phase column (Merck^®^) and a mobile phase composed of acetonitrile:methanol:isopropanol (85:10:5 v/v). The separation of carotenoids and chlorophyll pigments was carried out with a 1.5 mL/min flow rate, at room temperature and under isocratic conditions. The identification of the pigments was performed as described by (Simpson et al., 2016).

### RNA Extraction and Quantitative RT-PCR

RNA extraction was obtained from 20-100 mg of fresh carrot tap roots or leaves and entire 2-weeks-old Arabidopsis plants that were pulverized with liquid nitrogen, homogenized with CTAB buffer (2% (w/v) CTAB, 2% (w/v) PVP40, 25 mM EDTA, 2 M NaCl, 100 mM Tris-HCl (pH 8.0) and 0.05 % spermidine trihydrochloride) and precipitated with LiCl (10 M) overnight. RNA was resuspended in nuclease-free water and the genomic DNA traces were eliminated by a 40 min DNAse I treatment. For cDNA synthesis, 3 μg of total RNA was incubated with 1mM of Oligo-AP primer (Supplementary Table I) and Improm II reverse transcriptase (Promega^®^) according to the manufacturer’s recommendations. Quantitative RT-PCR (qRT) was performed as described in Fuentes et al. (2012) and Moreno et al. (2013) in a Stratagene Mx3000P thermocycler, using SYBR Green double strand DNA binding dye. AtqPAR1.F and AtqPAR1.R primers were used to amplify a specific fragment of the coding sequences of *AtPAR1* (AT2G42870) and DcqPAR1.F and DcqPAR1.R for *DcPAR1* (XM_017390696.1) (Supplementary Table I). DcqUbi.F and DcqUbi.R primers were used to amplify *Ubiquitin* and AtqPP2A.F and AtqPP2A.R to amplify *PP2A* housekeeping genes. Specific primers of carotenogenic genes *DcPSY1*, *DcPSY2*, *DcLCYB2*, *DcLCYE*, *DcCHXB1*, *DcCHXB2* and *AtPSY* were the same as described in Fuentes et al. (2012) and summarized in Supplementary Table I. The qRT analysis was performed with three biological replicates and two technical repeats and all reaction specificities were tested with melting gradient dissociation curves. To test for significant differences in gene expression, results were subjected to a one-way Anova (p<0.05, confidence interval 95%) with Dunnet post-test according to the General Linear Models option in the statistical software package Graphpad Prism. The relative transcript levels were obtained using the PfaffI equation (Pfaffl, 2001).

### Bimolecular fluorescence complementation (BiFC) Assay

The PCR8:DcPAR1 vector was recombined with BiFC vectors (Gateway^™^-based BifC binary vectors, pYFC43 and pYFN43; Belda-Palazón et al., 2012), obtaining N-YFP:DcPAR1 and C-YFP:DcPAR1. The BiFC binary vectors (Belda-Palazón et al., 2012) and N-YFP:AtPIF7 and C-YFP:AtPIF7 vectors were provided by Dr. Jaume Martínez-García (IBMCP CSIC-UPV, Valencia, Spain). The four vectors were transformed in *Agrobacterium* GV3101 and used for *Nicotiana benthamiana* agroinfiltration (Simpson et al., 2018). Four days after injection, YFP fluorescence was detected under a LEICA TCS SP5 confocal microscope with a light excitation wavelength of 488 nm and a filter for YFP at 520-560 nm. The images were obtained and processed with Leica LAS AF Lite software.

### Arabidopsis protein extraction and western blot

The Arabidopsis protein extraction was performed according to Saucedo et al. (2019) with modifications. Briefly, 400 mg of fresh tissue of entire 2-weeks-old Arabidopsis plants were pulverized with liquid nitrogen and then 4 ml of extraction buffer [0.5 M TrisHCl pH8, 0.7 M Sucrose, 2 protease inhibitor tablets (complete Mini, EDTA-free-Thermo Scientific Lot # UG27666820), 50 mM EDTA, KCl 0.1 M, β-mercaptoethanol 0.2%)] were added and homogenized until the sample was completely thawed. Then, 2 ml of basic saturated phenol pH 8.0 (Winkler # 20192088) were added, shaken 10 minutes in ice and centrifuged at 8000 g for 19 min at 4 °C. The supernatant was recovered and four volumes of 0.1 M ammonium acetate in methanol (Merck gradient for liquid chromatography) were added, and proteins were precipitate overnight at −20 °C. After centrifugation the pellet was washed with 0.1 M ammonium acetate in methanol, and with acetone. The remaining acetone was evaporated at room temperature and the proteins (pellet) were stored at −20 °C in resuspension buffer (100 mM Tris-HCl pH 7.0, 1% SDS). The extraction was carried out in triplicate for each transgenic line and control.

